# Social Behavior in Mice Relies on a Capacity to Phosphorylate the Serotonin Transporter

**DOI:** 10.64898/2026.11.23.768675

**Authors:** Carina Arnold, Adele Stewart, Hideki Iwamoto, Angelina Emerson, Tristan C. Wells, Manuela G. Zapata, Sammanda Ramamoorthy, Randy D. Blakely

## Abstract

The biogenic amine 5-hydroxytryptamine (serotonin, 5-HT) regulates social behavior across the animal kingdom. Although drugs that promote 5-HT transporter (SERT)-mediated 5-HT efflux within the nucleus accumbens (NAc) enhance sociability, the extent to which endogenous mechanisms regulating SERT contribute to normal social behavior is unknown. Prior *in vitro* studies have shown that SERT trafficking and activity are controlled by transporter phosphorylation, with the Thr276 site required for rapid, protein kinase G (PKG)-dependent increases in SERT function. Using SERT Ala276 knock-in mice, we demonstrate that sex-specific social behaviors rely on the capacity to phosphorylate SERT at Thr276. Whereas WT males and females exhibited social preference and sustained interactions with juvenile mice, these behaviors were selectively diminished in Ala276 females. In contrast, Ala276 males exhibited preserved sociability but increased retreat behavior in the tube dominance test. Dorsal and ventral brain slice biotinylation studies revealed sex-independent, PKG-induced increases in striatal SERT surface expression that were abolished in Ala276 mice. In the dorsal raphe nucleus (DRN), PKG activation failed to elevate SERT surface expression in either sex. Extracellular 5-HT responses in the NAc recorded via *in vivo* fiber photometry during repeated, but not initial, social interactions were absent in Ala276 females but not males. *Ex vivo* patch clamp recordings from DRN slices uncovered reduced excitability of 5-HT neurons from female Ala276 mice compared with WT controls, an effect reversed by blockade of 5-HT_1A_ autoreceptors, consistent with tonic activation of these receptors in mutants. Supporting this idea, systemic administration of the 5-HT_1A_ partial agonist buspirone restored sociability in Ala276 females and social dominance in Ala276 males, while elevating extracellular NAc 5-HT responses in Ala276 females. Our findings support a model in which sustained social behavior requires the capacity to phosphorylate SERT at Thr276 to limit somatodendritic 5-HT_1A_ autoreceptor-mediated inhibition and support serotonergic signaling.

## INTRODUCTION

Social behavior is an evolutionarily conserved process essential for survival and reproductive success [1, 2]. Across species, social behaviors support bonding, nurturing and mate selection, the persistence of supportive social interactions, and the regulation of aggression and dominance hierarchies. These functions rely on neural systems that assign motivational value to social stimuli, enabling organisms to seek, maintain, or avoid social contact in a context-dependent manner [3, 4]. The monoamine neurotransmitter serotonin (5-HT) is an evolutionarily conserved modulator of social behavior [5]. Across species, 5-HT signaling influences social approach, affiliative interactions, and social reward [6-8]. In mammals, serotonergic neurons of the dorsal raphe nucleus (DRN) project throughout the forebrain including regions that dictate social cognition and motivation, including the hippocampus, prefrontal cortex, and ventral striatum (vStr), within which the nucleus accumbens (NAc) is particularly linked to social behavior[7, 8]. Within these circuits, 5-HT can regulate social behavior not only through postsynaptic receptor signaling[7, 9] but also through presynaptic and somatodendritic autoreceptor-mediated feedback which has been shown to limit serotonergic neuron excitability and 5-HT release[10-12].

Deficits in sociability are found in multiple neurobehavioral disorders including major depressive disorder (MDD)[13], anxiety disorders, and Autism Spectrum Disorder (ASD)[14]. Although selective 5-HT reuptake inhibitors (SSRIs), blockers of the 5-HT transporter (SERT), are among the most prescribed pharmacological treatments for mood disorders, 5-HT elevations triggered by these agents are not brain circuit specific nor temporally entrained to episodic behavioral states evoked by naturalistic social stimuli[8, 12]. These limitations underscore a need to better understand how 5-HT signaling, and its regulation by SERT in discrete pathways, contributes to behavioral disorders.

One of the most striking demonstrations of serotonergic control of social behavior comes from the prosocial effects of the psychostimulant (±)3,4-methylenedioxymethamphetamine (MDMA)[12]. MDMA displays prosocial effects across diverse species, ranging from invertebrates such as crayfish and octopi to rodents and humans[5, 12, 15, 16], highlighting a deeply conserved serotonergic mechanism supporting sociability. At a molecular level, MDMA triggers non-vesicular 5-HT release through reversal of SERT at the plasma membrane, including axonal membranes of serotonergic projections to the NAc[17]. Increased 5-HT signaling within the NAc medial shell is a powerful driver of social engagement[8]. Consistent with this idea, whereas SERT blockade promotes social interaction, animal models expressing an ASD-associated hyperfunctional SERT variant display social deficits[6, 18]. Together, these findings emphasize the role of SERT-dependent 5-HT release as a powerful driver of social engagement and position SERT as a major determinant of socially relevant 5-HT signaling.

SERT activity is dynamically regulated by intracellular signaling pathways that control transporter surface trafficking as well as the intrinsic rate of 5-HT transport[19, 20], the latter thought to arise from differential residence in rate-limiting conformational states[21, 22]. Multiple kinase cascades, including protein kinase C (PKC), p38 MAPK, GSK3β and cGMP-dependent protein kinase (PKG), regulate SERT trafficking and function[23-25]. Importantly, SERT coding variants found in multiple neurobehavioral disorders[26, 27] including ASD[28-30] disrupt these mechanisms. PKG signaling has been reported to regulate both trafficking-dependent and -independent modes of SERT-mediated 5-HT uptake and clearance *in vitro* and *ex vivo*[31, 32]. Systematic mutational analysis of SERT identified threonine 276 (Thr276), within the second intracellular loop, as uniquely required for PKG-dependent SERT phosphorylation and stimulation of 5-HT transport[32]. Molecular modeling together with structural analyses of SERT captured in inward- and outward-facing conformations indicate that Thr276 becomes accessible to cytosolic regulators when SERT is in an inward-facing state[22, 33]. In this conformation, phosphorylation at Thr276 is proposed to accelerate the return of the transporter to an outward-facing state, which is believed to be the rate-limiting step in the 5-HT transport cycle[34, 35].

To directly assess the *in vivo* role of PKG-dependent SERT phosphoregulation in 5-HT-regulated behavior, we generated Ala276 knock-in (KI) mice, rendering the transporter unable to be phosphorylated by PKG while preserving basal transporter expression and 5-HT uptake capacity[34]. Here, using a combination of *in vivo* and *ex vivo* recording approaches, we show that SERT Ala276 mice demonstrate sex-dependent alterations in 5-HT autoreceptor signaling and sociability, establishing SERT phosphoregulation as a potential target for novel therapeutics, particularly for disorders that display perturbations in social behavior.

## RESULTS

### Social preference is disrupted in a sex-dependent manner by the Ala276 mutation

We have previously reported sex-dependent behavioral alterations in Ala276 mice, including increased losses in the tube test in males[34], often interpreted as a decrease in social dominance. To directly assess social preference in a paradigm that does not constrain movement or enforce interaction, we performed the three-chamber social preference test, which allows mice to more freely interact with either a sex- and age-matched novel mouse or a novel object. No sex or genotype effect was observed with respect to overall exploration of the testing chamber (Fig. 1A). Similarly, both male and female Ala276 mice clearly demonstrated social preference (Fig. 1B). However, when relative sociability[12], was quantified as the normalized preference for the social stimulus over the object, Ala276 females selectively exhibited a significantly reduced preference for the novel mouse-containing zone (Fig. 1C), as well as reduced direct sniffing compared with WT littermates (Fig. 1D). These findings suggest that although the Ala276 mutation does not abolish social preference, it selectively diminishes social interest in females. To determine whether this reduction in social interest also affects direct social engagement, we next assessed social interactions using a juvenile interaction assay conducted with freely moving mice. In this less constrained paradigm, we found that male Ala276 mice demonstrated no reduction in social interactions relative to their WT littermates (Fig. 1E-G). Female Ala276 mice, in contrast, showed clear differences in this assay. Specifically, they exhibited a significant reduction in the number of interaction bouts across the full 5-min test period (Fig. 1E). Because each interaction bout reflects an independent initiation of social contact following disengagement, bout number provides a measure of an animal’s propensity to repeatedly re-engage with a social partner over time. Active interaction time also showed a trend toward reduction in Ala276 females but did not reach statistical significance (*p* = 0.065; Fig. 1F). These findings suggest that the primary social deficit in female Ala276 mice lies in repeatedly reinitiating social interactions rather than in the duration of individual interactions, once initiated. To assess the proportion of social interactions that were active, we calculated interaction ratios (see Methods) for each genotype. We found that Ala276 females exhibited a significantly reduced social interaction ratio compared with WT littermates (p < 0.01; Fig. 1G), with two-way ANOVA revealing a significant sex × genotype interaction (*p* = 0.02). Together, these findings demonstrate that the inability to phosphorylate SERT at Thr276 produces a female-specific reduction in sociability, characterized by preserved social preference but a diminished propensity to repeatedly engage with a social partner.

**Figure 1.**
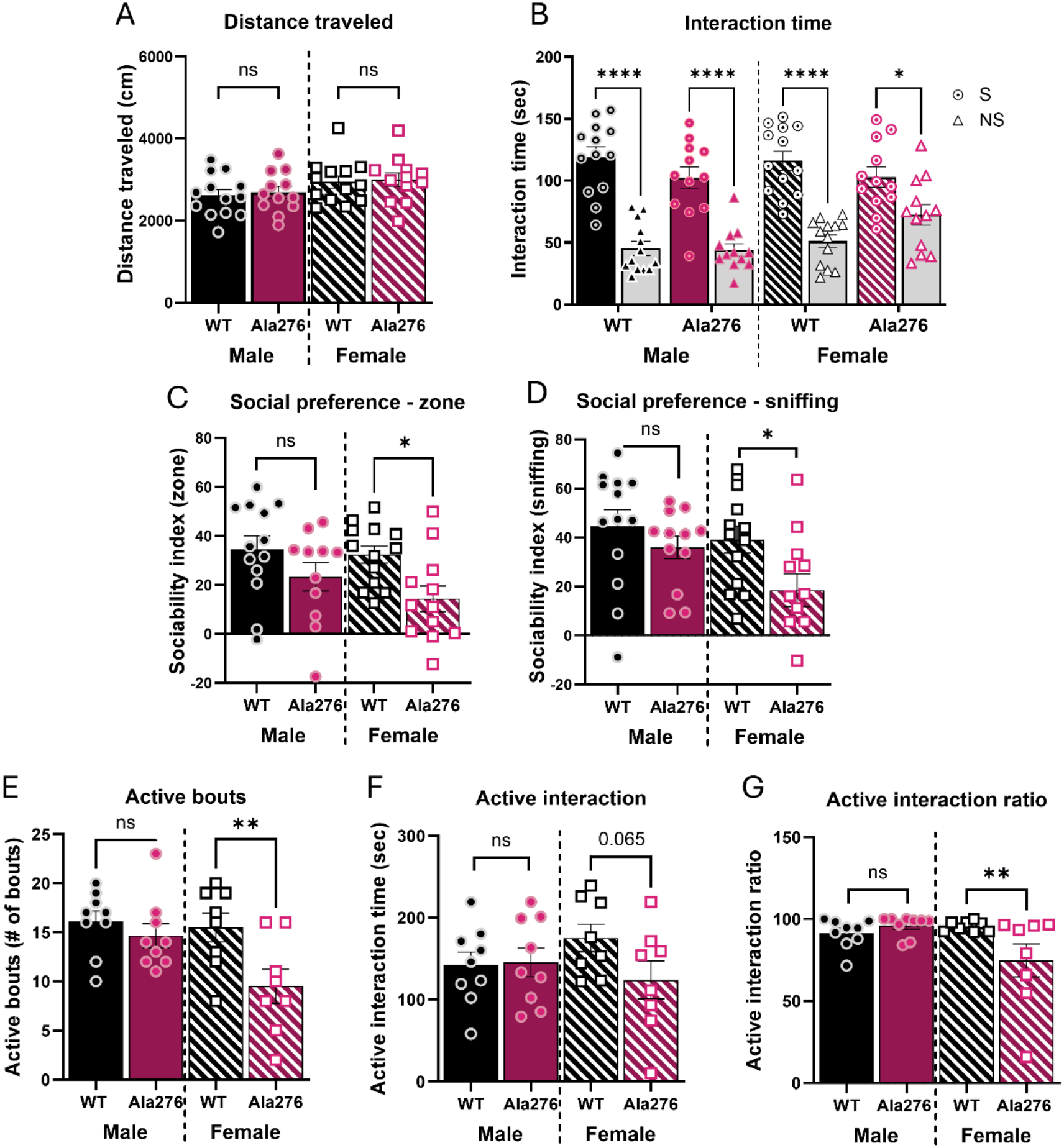
Ala276 females display reduced sociability in the three-chamber social preference test. (**A**) Total distance traveled during social preference test comparing male (left) and female (right) WT and Ala276 mice (n = 12–13 per group). (**B**) Time spent interacting with a novel social stimulus mouse (S) versus a non-social, inanimate object (NS) during a 10 min test. (**C–D**) Sociability index quantified from 10 min of exploration of the chamber zone (**C**) and defined sniffing zone (**D**) around the enclosures. (**E–G**) Behavioral measures from the juvenile interaction approach. (**E**) Number of active interaction bouts initiated by the experimental mouse, (**F**) active interaction time, and (**G**) social interaction ratio, comparing male WT (n = 9) and Ala276 (n = 9) mice on the left and female WT (n = 9) and Ala276 (n = 7) mice on the right. Data are presented as mean ± SEM. Statistical significance for interaction with the social versus non-social stimulus was determined using multiple t-tests. Genotype and sex effects were analyzed using two-way ANOVA followed by uncorrected Fisher’s LSD multiple comparisons test. ns, not significant; *p < 0.05, **p < 0.01, ****p < 0.0001. In **G**, a significant sex × genotype interaction was detected for the social interaction ratio (p = 0.020).

### SERT Ala276 does not impact anxiety-related behavior

Because serotonergic perturbations are frequently linked to anxiety, we next asked whether a generalized anxiety-like phenotype could account for the sex-dependent social deficits observed in Ala276 mice. Consistent with our prior report[34], Ala276 KI mice spent comparable time in the center versus periphery of the open field relative to WT littermates, and also showed no differences in the elevated plus maze, where increased time in enclosed arms is typically interpreted as anxiety-like behavior. To support this interpretation in an independent assay, we employed the light/dark test and again observed no genotype differences in time spent in the light compartment in either sex (Fig. S3A-F). Collectively, these null results support the conclusion that the social deficits in Ala276 mice are not secondary to heightened anxiety or altered avoidance behavior, but more likely reflect a selective disruption of 5-HT-modulated social circuitry.

### A capacity to phosphorylate Thr276 is required for PKG-dependent SERT trafficking in serotonergic projections

Having identified sex-dependent social deficits in Ala276 mice, we next asked how loss of the capacity to phosphorylate Thr276 alters SERT regulation in intact brain slices. SERT is regulated through changes in catalytic activity and/or surface trafficking, which *in vitro* studies indicate can occur in series or independently[31, 36]. Multiple studies from our group have demonstrated that activation of PKG increases SERT surface expression in heterologous systems [19, 30, 31].

Ramamoorthy and colleagues found that PKG activation by 8-Br-cGMP increased SERT activity in rat midbrain synaptosomes without a change in surface expression[32]. We sought to extend these findings to intact brain preparations, where endogenous circuitry, cellular architecture, and neuromodulatory inputs are preserved. Thus, we asked whether 1) the loss of a capacity for phosphorylation at Thr276 alters basal SERT surface expression and/or 2) precludes the ability of a PKG activator to further elevate SERT surface expression *ex vivo* using acute brain slices from WT and Ala276 mice. We prepared midbrain slices to evaluate basal and PKG-dependent regulation of SERT surface expression in an area enriched for serotonergic cell bodies. We also prepared striatal slices from male and female mice, subdivided into dorsal and ventral regions due to their distinct functional roles, with the latter region containing DRN^5-HT^ projections that contribute to social behavior [6, 8, 12]. SERT surface expression was quantified following affinity purification of proteins covalently tagged with the membrane-impermeant biotinylation reagent sulfo-NHS-SS-biotin[37]. After acute treatment with vehicle or the PKG activator 8-bromo-cGMP (8-Br-cGMP, 250µM, 30min) surface labeled SERT proteins were isolated and quantified as described in Methods. We first examined SERT surface expression in the DRN, where alterations in transporter availability could influence local serotonergic signaling. Reduced basal surface SERT levels were detected in male Ala276 DRN compared with WT, a difference that was not evident in females (Fig. 2A,D). Notably, 8-Br-cGMP treatment of these preparations had no effect on SERT surface expression of either genotype or sex (Fig. 2A,D). These findings indicate that PKG-dependent regulation of SERT in the DRN is unlikely to involve changes in transporter surface trafficking, consistent with trafficking-independent changes in SERT activity observed in midbrain synaptosomes[32].

**Figure 2.**
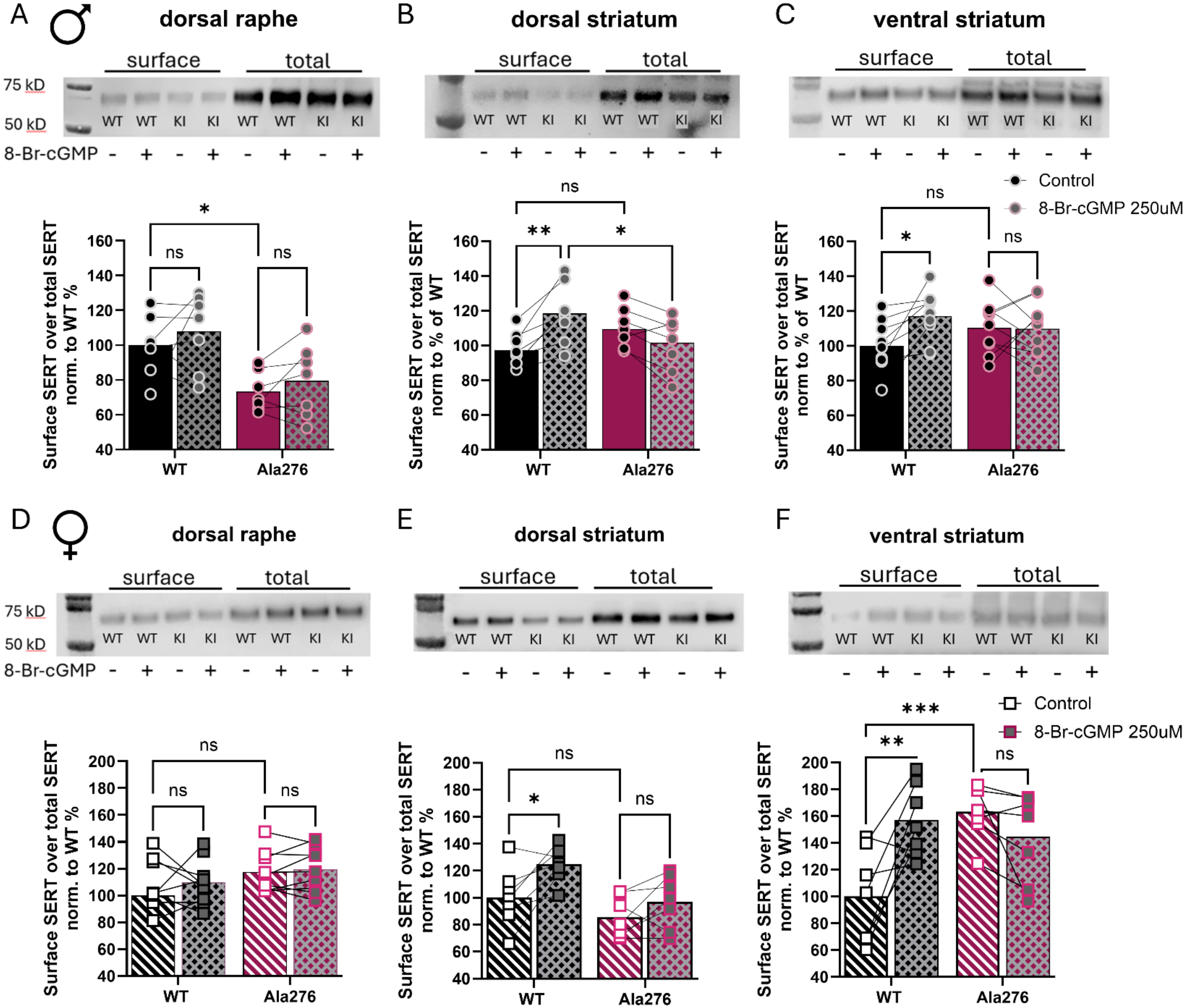
SERT Ala276 mice lack 8-Br-cGMP induced elevations in SERT surface expression in striatal projections. (**A-F**) SERT surface expression in DRN (**A**,**D**), dorsal striatum (**B**,**E**), and ventral striatum (**C**,**F**) of WT and Ala276 male and female mice (n = 8 per group). Acute brain slices were hemisected and incubated for 30 min with vehicle (-) or 250 µM 8-bromo-cGMP (+). Data are presented as mean ± SEM. Statistical analysis was performed using two-way ANOVA followed by Šídák’s multiple comparisons test. ns, not significant; *p < 0.05, **p < 0.01, ***p < 0.001

Given the absence of PKG-dependent SERT trafficking in the DRN, we next examined whether this regulation differs in striatal projection regions, particularly the ventral striatum, where serotonergic signaling contributes to social behavior[12]. Biotinylation studies with striatal slices from males revealed no alteration in basal surface SERT expression in dorsal or ventral preparations comparing Ala276 and WT (Fig. 2B,C). Female dorsal striatal slices also showed no basal effects of genotype on surface expression (Fig. 2E). Interestingly, basal SERT surface expression was elevated in the ventral striatum of female Ala276 mice compared with WT (Fig. 2F). In contrast with our findings in DRN preparations, 8-Br-cGMP treatments significantly increased SERT surface expression in WT slices prepared from both striatal regions in a sex-independent manner (Fig. 2B,C,E,F). Notably, this regulation was absent in Ala276 slices from both sexes. Together, these findings demonstrate that PKG-dependent SERT surface trafficking differs between the DRN and striatal projection regions and requires the capacity to phosphorylate Thr276 in the latter.

### Real-time monitoring of extracellular 5-HT dynamics during social encounters reveals impaired NAc 5-HT signaling in female Ala276 mice

Having established that loss of a capacity to phosphorylate Thr276 disrupts female social behavior and dynamic regulation of SERT trafficking in regions containing serotonergic projections, including the vSTR where 5-HT is known to govern social behavior[38], we next asked whether this altered regulation is accompanied by changes in extracellular 5-HT dynamics during social interactions. We hypothesized that the reduced sociability of Ala276 females may be associated with altered NAc 5-HT signaling. To pursue this hypothesis, we expressed a genetically encoded 5-HT sensor (GRAB_5-HT3.6_) in the medial shell of the NAc of WT and Ala276 mice (Fig. S1A) along with an optical probe for fiber photometry[8]. Three weeks following surgery, we monitored non-vesicular 5-HT release in response to 7.5 mg/kg MDMA i.p. as a functional test of sensor expression. We detected a robust and comparable increase in extracellular 5-HT after MDMA treatment of either WT or Ala276 mice irrespective of subject sex (Fig. S1C,D), consistent with no effect of the Ala276 mutation or sex on MDMA potency, as assessed *ex vivo* using synaptosomal [^3^H]5-HT uptake assays (Fig S1F-M). Having established comparable sensor responses and MDMA sensitivity across genotypes, we next recorded GRAB_5-HT_ fluorescence during social encounters using the juvenile interaction assay, a test previously shown to reveal correlations between NAc 5-HT levels and mouse social behavior[8]. In males, we failed to detect significant changes in social stimulus-dependent 5-HT signals during the first interaction, comparing Ala276 to WT controls (Fig. 3A). Although a second exposure of male Ala276 mice to the juvenile stimulus resulted in a trend toward a reduction in extracellular 5-HT signaling relative to WT controls (Fig. 3B), this effect did not reach statistical significance. As seen with males, WT and Ala276 females exhibited a comparable increase in NAc 5-HT signals on the initial social interaction (Fig. 3C). In striking contrast, during the second interaction, the social stimulus-induced 5-HT response was absent in Ala276 females, whereas WT females maintained a clear elevation in extracellular 5-HT (Fig. 3D). Thus, the loss of Thr276 phosphorylation does not prevent the initial serotonergic response to a social stimulus but selectively impairs its recurrence during subsequent social engagement in females. To directly compare serotonergic dynamics between sexes, we replotted the fiber photometry responses to overlay male and female traces. As shown in Fig. 3E, males displayed a delayed onset of the 5-HT signal during the first interaction, followed by a rapid rise in signal, whereas females exhibited a more gradual, sustained increase from the onset of interaction. Despite these initial temporal differences, however, analysis of these profiles across the recording period revealed no overall sex-dependence of 5-HT elevations. Similarly, during the second interaction, WT males and females showed an equivalent increase in extracellular 5-HT (Fig. 3F). Ala276 males and females exhibited similar 5-HT responses during the first interaction (Fig. 3G), paralleling the response profiles of WT males and females. During the second interaction, however, Ala276 males displayed a modest increase in extracellular 5-HT that was significantly larger than the response in Ala276 females (Fig. 3H), whose average signal fell below the pre-interaction baseline. Together, these findings reveal a sex-dependent disruption of NAc serotonergic signaling in Ala276 mice that emerges during repeated, rather than initial, social encounters.

**Figure 3.**
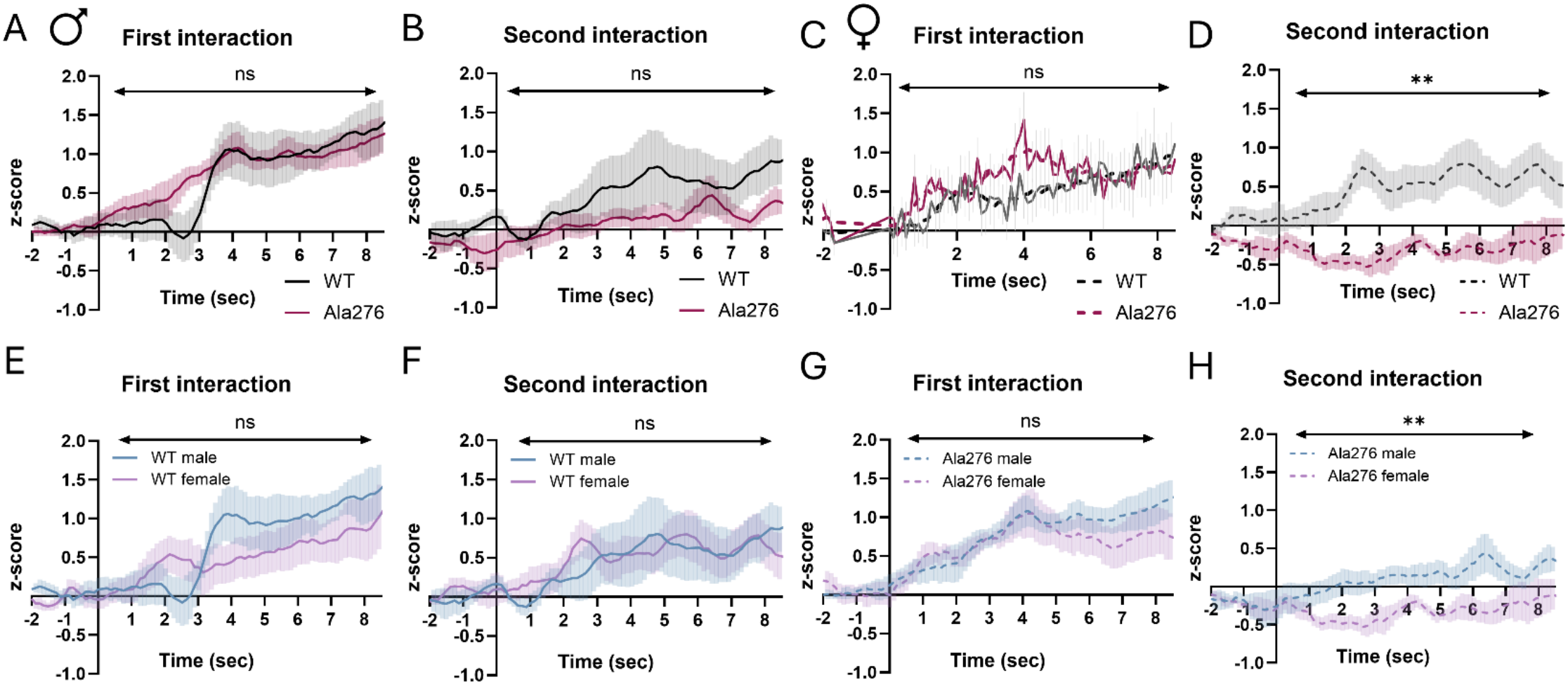
Loss of social stimulus-dependent 5-HT elevations in the NAc of Ala276 females. Male **(A, B)** GRAB_5-HT_ fluorescence traces during the first **(A)** and second **(B)** juvenile interaction, comparing WT (*n* = 7) and Ala276 (*n* = 9) males. Female **(C, D)** GRAB_5-HT_ fluorescence traces during the first **(C)** and second **(D)** juvenile interaction, comparing WT (*n* = 7) and Ala276 (*n* = 7) females. **(E, F)** GRAB_5-HT_ fluorescence traces during the first **(E)** and second **(F)** juvenile interaction, comparing WT males (*n* = 7) and females (*n* = 7). **(G, H)** GRAB_5-HT_ fluorescence traces during the first **(G)** and second **(H)** juvenile interaction, comparing Ala276 males (*n* = 9) and females (*n* = 7). Data are presented as mean ± SEM. Statistical comparisons were performed using two-way RM-ANOVA. ns, not significant; **p < 0.01

### Enhanced 5-HT_1A_ autoreceptor-mediated inhibition suppresses the firing of DRN^5-HT^ neurons in brain slices from Ala276 females

The failure of Ala276 females to sustain NAc 5-HT signaling during repeated social encounters raised the possibility that diminished serotonergic output originates upstream at the level of DRN^5-HT^ neuron activity. We therefore asked whether loss of the capacity to phosphorylate SERT at Thr276 alters DRN^5-HT^ neuron firing. Because the deficit in sustained NAc 5-HT signaling was most pronounced in females, we focused these studies on female mice. We monitored the firing properties of DRN^5-HT^ neurons using whole-cell recordings under current-clamp in acute brain slices. Neurons were activated with phenylephrine (10 µM) to induce a regular tonic firing pattern via alpha1-adrenergic stimulation (1–3 Hz), mimicking physiological drive of 5-HT neurons during wakefulness[10]. Under these conditions, DRN^5-HT^ neurons from Ala276 females exhibited a significantly reduced firing rate compared with WT controls (Fig. 4A,B). To determine whether this reduction in firing reflects enhanced inhibitory feedback, we next examined the contribution of 5-HT_1A_ autoreceptor signaling. DRN^5-HT^ neuron excitability is known to be tightly regulated by inhibitory 5-HT_1A_ autoreceptors with SERT-mediated 5-HT clearance serving as major negative regulator of 5-HT_1A_ receptor activation[39]. We thus asked whether enhanced tonic autoreceptor-mediated feedback, as might be the case with a SERT unable to efficiently transition to an activated state due to lack of Thr276-SERT phosphorylation, contributes to the basal reduction in firing observed in our Ala276 slices. At the firing rate we imposed on DRN^5-HT^ neurons with phenylephrine (1-3Hz), the 5-HT_1A_ antagonist WAY-100635 did not elevate firing rate in WT neurons, consistent with sufficient SERT-mediated 5-HT clearance to prevent 5-HT_1A_ autoreceptor-mediated inhibition (Fig. 4A,B). In contrast, in recordings from Ala276 mice, WAY-100635 elevated the firing rate of DRN^5-HT^ neurons, achieving a rate comparable to that observed in WT neurons. Together, these findings indicate that enhanced 5-HT_1A_ autoreceptor-mediated inhibition contributes to the reduced firing of DRN^5-HT^ neurons in Ala276 females, an effect that can be reversed by pharmacological blockade of these receptors.

**Figure 4.**
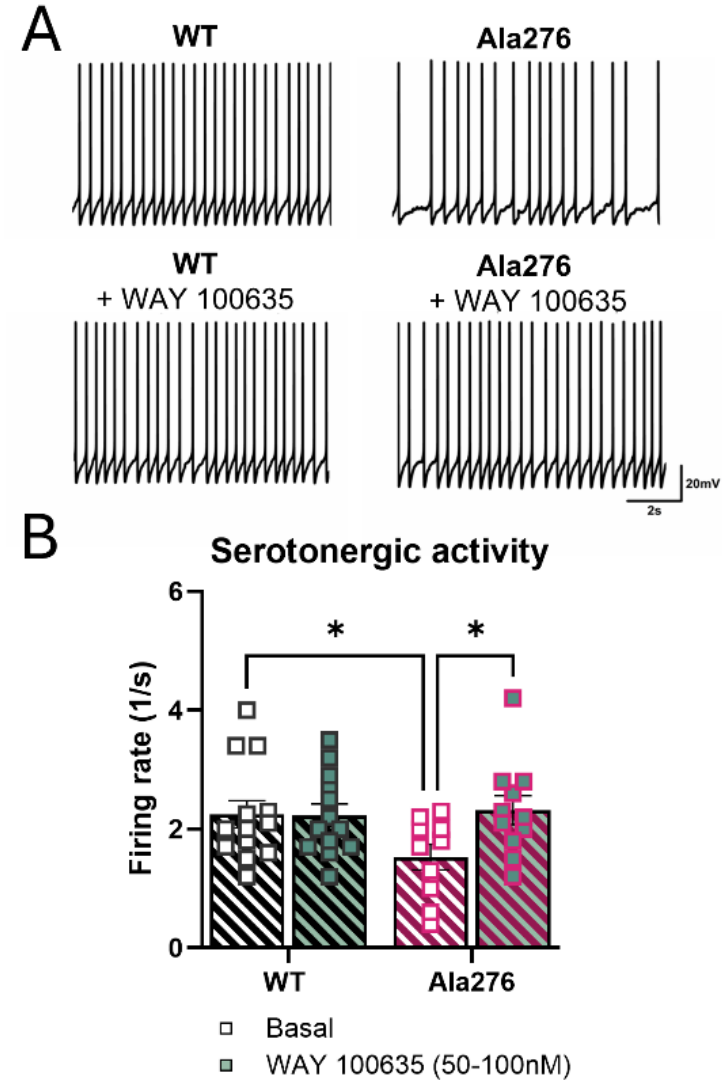
5-HT_1A_ receptor modulation rescues reduced DRN firing in female Ala276 mice. **(A)** Representative traces of DRN 5-HT neurons from WT (left) and Ala276 females (right) in response to (10 µM) phenylephrine under basal conditions (top) and following application of the (50-100 nM) 5-HT_1A_ receptor antagonist WAY-100635 (bottom). **(B)** Quantitative representation of neuronal activity with each point representing the firing rate of a single neuron. Data were obtained from WT (n = 5), KI (n = 6), WT + WAY-100635 (n = 5), and KI + WAY-100635 (n = 6) and are shown as mean ± SEM. Statistical analyses were performed using two-way ANOVA followed by Šídák’s multiple comparisons test, ns, not significant; *p < 0.05.

**Figure 4.**
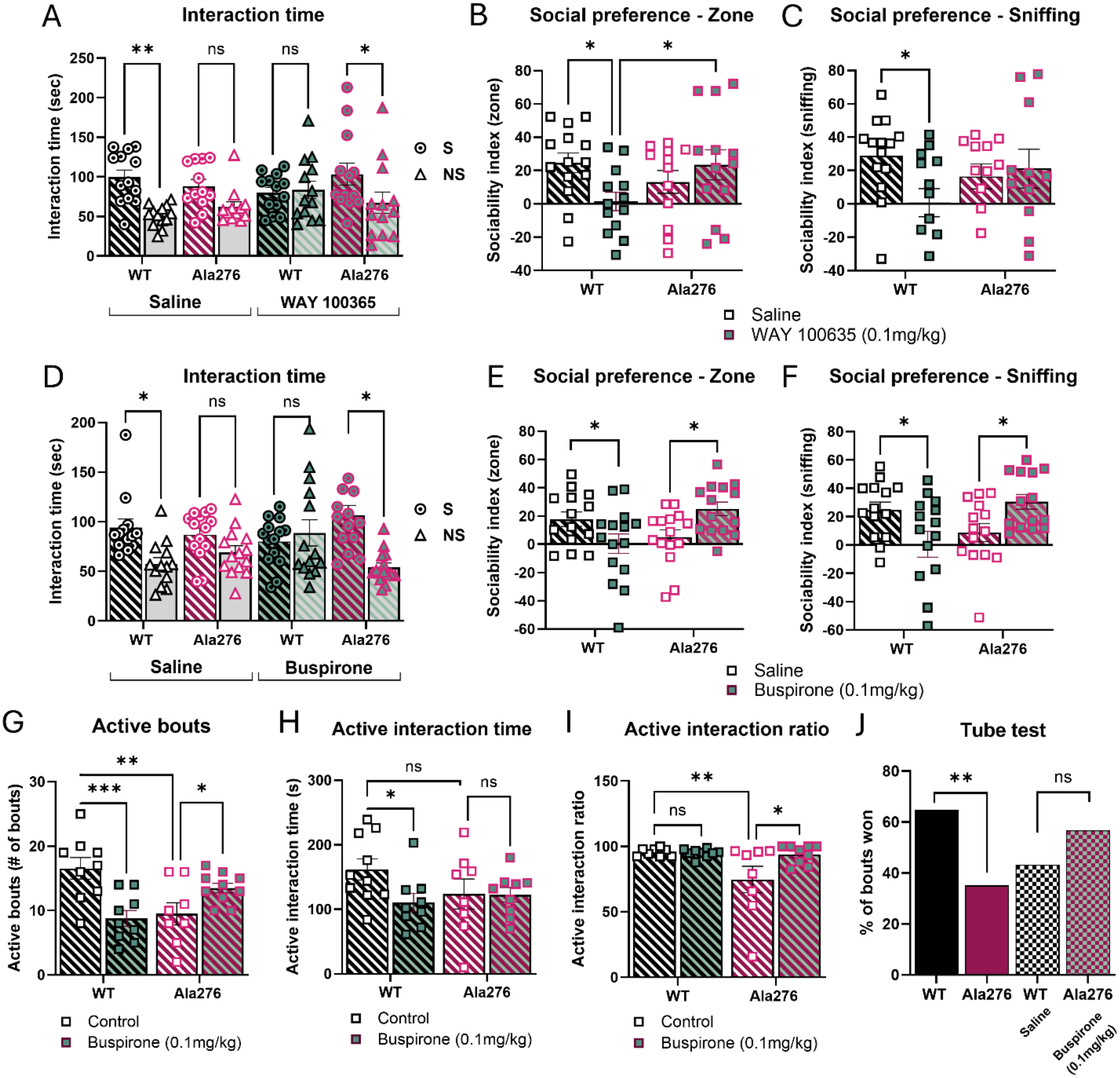
5-HT_1A_ receptor modulation rescues sociability deficits in Ala276. (**A-C**) Three-chamber social preference test following pretreatment with saline (WT n=14, Ala276 n=12) or WAY-100635 (0.1 mg/kg, s.c., 10 min before test begin) (WT n=13, Ala276 n=13). (**A**) Time spent interacting with the social versus non-social stimulus. (**B**) Time spent in the social chamber zone. (**C**) Time spent in the defined sniffing zone. WAY-100635 reduced sociability in WT but not Ala276 females. (**D-F**) Social preference test following pretreatment with saline (WT n=14, Ala276 n=15) or the partial 5-HT_1A_ receptor agonist buspirone (0.1 mg/kg, i.p., 30 min before test begin) (WT n=15, Ala276 n=14). (**D**) Time spent interacting with the social stimulus mouse (S) versus the inanimate object (NS). (**E**) Time spent in the social chamber zone. (**F**) Time spent in the defined sniffing zone. (**G**) Number of active interaction bouts of WT and Ala276 females with (WT n=9, Ala276 n=10) and without pretreatment (WT n=9, Ala276 n=8) of buspirone. (**H**) Active interaction time. (**I**) Social interaction ratio. (**J**) Tube test in male WT (n=8) and Ala276 (n=16) mice following saline or buspirone (0.1 mg/kg, i.p., 40 min before testing). Data are shown as mean ± SEM. Statistical analyses were performed using two-way ANOVA followed by Šídák’s multiple comparisons test or Fisher’s LSD (G/H/I), ns, not significant; *p < 0.05, **p < 0.01, ***p < 0.001.

### Social deficits in Ala276 females are rescued by the 5-HT_1A_ partial agonist buspirone

Having found that the failure to sustain NAc 5-HT signaling in Ala276 females is accompanied by 5-HT_1A_ autoreceptor-dependent suppression of DRN^5-HT^ excitability, we next asked whether pharmacological manipulation of 5-HT_1A_ receptors could restore social behavior. Systemic administration of the full 5-HT_1A_ antagonist WAY-100635 only partially rescued sociability in Ala276 females (Fig. 5A-C), potentially reflecting the broad pre- and postsynaptic distribution of 5-HT_1A_ receptors[40]. We therefore turned to the partial agonist buspirone, which has been shown to exhibit preferential efficacy *in vivo* at DRN autoreceptors relative to postsynaptic 5-HT_1A_ receptors in hippocampus and forebrain [41-43]. Importantly, partial agonists can functionally antagonize receptor signaling under conditions of elevated endogenous agonist activity, raising the possibility that buspirone could counteract excessive somatodendritic 5-HT_1A_ autoreceptor activation in Ala276 females. We therefore administered a low dose of buspirone (0.1 mg/kg) 30 min before testing as previously described[44, 45]. As shown in Fig 5D, WT females displayed a significant preference for the social stimulus versus the nonsocial stimulus, whereas Ala276 females exhibited no preference. Interestingly, buspirone produced opposing effects on social preference depending on genotype. Buspirone treated WT females lost social preference, consistent with agonism of 5-HT_1A_ autoreceptors in the context of efficient somatodendritic 5-HT clearance. In contrast, buspirone restored social preference in Ala276 females (Fig. 5D), consistent with our hypothesis that excessive 5-HT_1A_ autoreceptor-mediated inhibition contributes to their social deficits. Similarly, whereas buspirone reduced preference for the social zone (Fig. 5E) and sniffing of the novel mouse (Fig. 5F) in WT females, deficits in these measures were completely eliminated by buspirone treatment of Ala276 females. To determine whether these genotype-dependent effects of buspirone extend to interactions monitored in freely moving animals, we examined the drug’s effects in the juvenile interaction assay (Fig 5G-I). Consistent with our findings with the three-chamber test paradigm, buspirone significantly diminished the number of social interaction bouts of WT females compared with saline controls (Fig. 5G). In contrast, buspirone significantly increased the number of interaction bouts in Ala276 females (Fig. 5G). Whereas buspirone also reduced active social interaction time in WT females, active interaction time in Ala276 females did not show a corresponding increase despite the restoration of interaction bouts (Fig. 5H). This distinction suggests that buspirone may preferentially restore the propensity to initiate social contact rather than the overall duration of active social engagement. Quantification of the social interaction ratio revealed no effect of buspirone on WT females, but a significant increase in buspirone-treated Ala276 females (Fig. 5I), indicative of a greater proportion of active versus passive social engagement in mutant females. Importantly, active interaction time reflects the absolute duration of social engagement initiated by the experimental mouse, whereas the interaction ratio expresses this active engagement relative to total interaction time, including interactions initiated by the juvenile. Thus, the differential effects of buspirone on active interaction time and interaction ratio suggest that distinct components of social behavior, including the initiation and maintenance of social interactions, may be differentially sensitive to 5-HT_1A_ receptor modulation and associated changes in serotonergic signaling.

**Figure 5.**
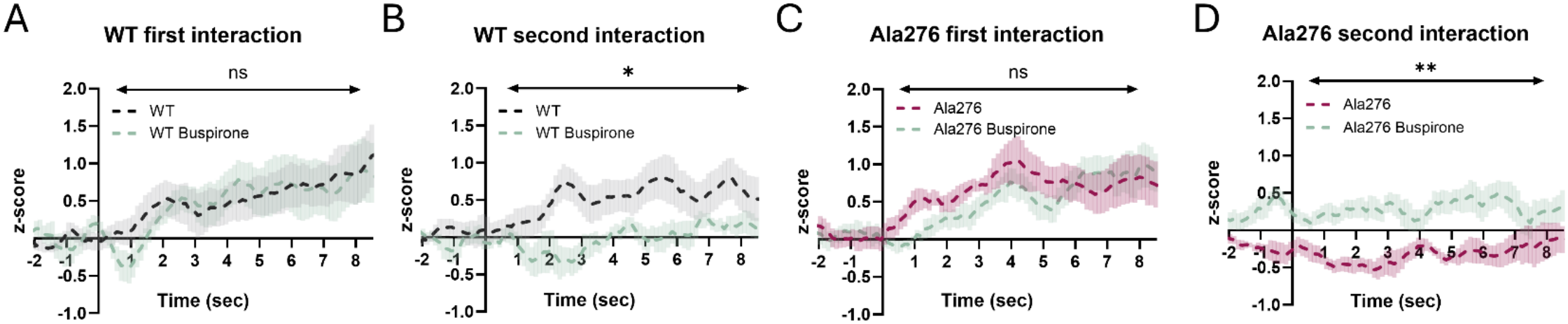
Buspirone restores NAc 5-HT signaling in Ala276 females. Fiber photometry recordings from WT females. (**A**,**B**) GRAB_5-HT_ fluorescence traces during the first (**A**) and second interaction (**B**) with a juvenile mouse comparing WT naïve (n=7) and WT buspirone injected (n=8) females. (**C**,**D**) GRAB_5-HT_ fluorescence traces during the first (**C**) and second interaction (**D**) with a juvenile mouse Ala276 naïve (n=7) and Ala276 buspirone injected (n=9) females. Data are presented as mean ± SEM. Statistical comparisons were performed using two-way RM-ANOVA. ns, not significant; *p < 0.05, **p < 0.01

Having demonstrated that buspirone restores several measures of social engagement in Ala276 females, we next asked whether its effects extend to the male-specific deficits in social dominance previously observed in Ala276 mice. We therefore administered buspirone to Ala276 males prior to interactions with novel WT males in the tube test. In our prior report, we found that compared with WT males, Ala276 males demonstrated reduced wins in the tube test, more often backing out of the tube upon encountering another male[34], a finding replicated in the current study (Fig. 5J). Following buspirone administration (0.1 mg/kg, i.p., 40 min before testing), this deficit was normalized, with Ala276 males displaying win rates comparable to WT controls. Together, these findings demonstrate that buspirone can reverse distinct social behavioral deficits associated with the Ala276 mutation in both sexes, despite their different behavioral manifestations.

### Buspirone restores social approach and NAc 5-HT levels in Ala276 females

Finally, we asked whether the pharmacological rescue of sociability by buspirone is accompanied by normalization of social stimulus-evoked extracellular 5-HT signaling in the NAc. Fiber photometry recordings were performed in WT and Ala276 females injected with buspirone (0.1 mg/kg) 40 min prior to juvenile interaction, as in the behavioral studies (Fig. S1B). For comparison, we plotted traces from buspirone-treated females alongside recordings obtained previously from animals without buspirone treatment, as shown in Fig. 3. In WT mice, extracellular 5-HT signals following buspirone administration closely followed the traces of untreated WT females during the first interaction (Fig. 6A). However, during the second interaction, buspirone-treated WT mice failed to show the stimulus-dependent increase in extracellular 5-HT previously observed in controls (Fig. 6B), consistent with the reduction in social interactions observed following buspirone administration (Fig. 5). In Ala276 females, as with WT females, buspirone did not alter the extracellular 5-HT response during the first interaction (Fig. 6C). Strikingly, however, buspirone partially restored the extracellular 5-HT response during the second social encounter in Ala276 females, which was markedly diminished in the absence of treatment (Fig. 6D). Although this response appeared attenuated relative to untreated WT controls, it was significantly greater than that observed in untreated Ala276 females. Together, these findings demonstrate that the rescue of social behavior by buspirone in Ala276 females is accompanied by a partial restoration of NAc extracellular 5-HT signaling during repeated social encounters, supporting a functional link between 5-HT_1A_ receptor modulation, serotonergic dynamics, and social engagement.

## DISCUSSION

Neurotransmitter transporters are dynamically regulated by intracellular signaling pathways, including protein kinases and phosphatases, thereby allowing rapid tuning of neurotransmitter clearance in response to neuronal activity[25]. Although PKG-dependent phosphorylation of SERT at Thr276 has been shown to enhance transporter activity and surface expression *in vitro* [30-32], the physiological significance of this post-translational modification has not been addressed *in vivo*. Here, through studies of the Ala276 knock-in mouse, we demonstrate an essential role for Thr276 phosphorylation in sustaining 5-HT signaling in a social context. We provide evidence that these deficits arise from excessive 5-HT_1A_ autoreceptor activation that diminishes excitability of DRN^5-HT^ neurons. Our studies reveal a remarkable requirement for a capacity to phosphorylate SERT at a single site to sustain specific aspects of complex social behavior, highlighting the selective contribution of this regulatory mechanism to distinct behavioral domains governed by 5-HT signaling[46, 47]. The SERT Ala276 mouse provides a mechanistically precise model to investigate the physiological importance of the capacity to phosphorylate Thr276 while preserving overall SERT expression and basal transport function.

Initial characterization of this line demonstrated normal SERT mRNA and protein expression, basal 5-HT uptake kinetics, tissue monoamine levels, anxiety-like behavior, and antidepressant-sensitive stress-coping behaviors despite disruption of this regulatory site[34]. Instead, the predominant phenotypes were sex-dependent alterations in social and repetitive behaviors, suggesting that the capacity to phosphorylate Thr276 is dispensable for maintaining basal serotonergic homeostasis but becomes important when serotonergic circuits must dynamically adapt to behavioral demands. In assays of sociability, female Ala276 mice exhibited the more prominent social phenotype, with deficits particularly evident during juvenile interaction. An evolutionarily conserved contribution of 5-HT to social behavior is supported by studies showing that serotonergic manipulation alters retreat behavior during agonistic encounters in crustaceans [48] and that MAOA-deficient mice, which exhibit markedly elevated brain 5-HT, display enhanced male aggression[49]. In humans, a point mutation resulting in MAOA deficiency was similarly associated with impulsive aggression in affected males[50]. Together, these findings underscore the importance of serotonergic homeostasis in shaping social behavior and suggest that disruption of dynamic SERT regulation at Thr276 can produce sex- and context-dependent alterations in social responses.

With respect to SERT, multiple gain-of-function mutations have been identified in subjects with ASD, with the Gly56Ala (G56A) substitution being the most intensively studied. Interestingly, the elevated 5-HT uptake activity displayed by this variant fails to respond further to PKG activation[30], suggesting that the structural conformation of this variant is constitutively biased toward an outward-facing state, a hypothesis supported by biochemical and imaging studies[21]. Together, these findings imply disrupted PKG-dependent regulation of SERT in altered serotonergic function and raise the question of how loss of the PKG-sensitive Thr276 phosphorylation site affects dynamic SERT regulation in native tissue. In keeping with prior finding of a PKG-linked pathway supporting elevated SERT surface expression and 5-HT transport capacity in cell lines and transfected cells, we found that 8-Br-cGMP, in a sex-independent manner, robustly enhanced SERT surface expression in dorsal/ventral slices prepared from WT striatum. This increase was abolished in Ala276 mice of both sexes. Notably, 8-Br-cGMP regulation of SERT surface expression was not observed in midbrain slices containing DRN^5-HT^ neurons, suggesting that phosphorylation of Thr276 selectively enhances SERT surface trafficking in projection areas versus somatodendritic compartments, where PKG-dependent catalytic activation predominates [32].

We speculate that activation of PKG mobilizes SERT transport vesicles in the striatum but not midbrain due to enrichment in axons versus somatodendritic compartments of specialized endocytic/exocytic mechanisms involved in membrane protein trafficking. PKG-linked pathways are known to influence actin filament rearrangement and synaptic vesicle fusion and recycling[51-53], likely pertinent for other transport vesicles[54-56]. Such an idea is consistent with other reports showing 8-Br-cGMP increasing SERT activity, but not trafficking, in midbrain preparations[32]. Structurally, Thr276 resides within the conformationally dynamic intracellular loop 2 (IL2), becoming accessible during inward-facing transitions of the transport cycle[22, 34]. Phosphorylation of Thr276 may facilitate the rate-limiting return of SERT to an outward-facing conformation, thereby accelerating 5-HT transport independent of changes in surface expression. Interestingly, studies have shown that neuronal nitric oxide synthase (nNOS) retains SERT intracellularly via a direct physical interaction. Chanrion et al [57] speculate that activation of nNOS bound to SERT may acutely increase activity of surface SERTs in a trafficking-independent manner, which we suggest may derive from the ability of nitric oxide to increase cGMP levels, activate PKG, and catalytically stimulate the transporter via Thr276. Our fiber photometry findings identify impaired serotonergic signaling within the NAc as the circuit-level correlate of the female behavioral phenotype. 5-HT release in the NAc plays a critical role in sociability induced by MDMA[12] and social reward[8], with recent studies showing that increases in 5-HT during social interactions arise from direct DRN serotonergic inputs into the medial shell of the NAc[8]. Conversely, inhibition of DRN^5-HT^ neurons reduces extracellular 5-HT levels in the NAc and sociability[8], mimicking 5-HT deficits in autism models that exhibit reduced sociability[58].

In our studies, the initial NAc 5-HT response during the first interaction was equivalent between genotypes, indicating that disruption of the capacity to phosphorylate Thr276 does not impair the initiation of serotonergic signaling. Rather, the mutation appears to limit the ability to sustain serotonergic signaling that can maintain repeated social encounters. Importantly, genotype differences were absent in MDMA-induced 5-HT release in both sexes (Fig. S1C,D). Because MDMA induces non-vesicular, SERT mediated 5-HT efflux[59-61], it is unlikely that the intrinsic transport function of SERT is perturbed by the Ala276 mutation, as was previously reported in transfected cells[32]. Rather, the mutation reveals a requirement for phosphorylation at Thr276 to achieve dynamic changes in 5-HT signaling needed to sustain social engagement. One mechanism that could contribute to the loss of 5-HT signaling needed to support social encounters in the Ala276 mice is negative feedback regulation of vesicular release by terminal 5-HT_1B_ autoreceptors [62]. Deficits in turn limit activation of postsynaptic 5-HT_1B_ receptors in the NAc that have been shown to be essential for the prosocial actions of MDMA[12]. Interestingly, Ala276 females exhibit elevated NAc SERT surface expression, approaching levels observed in WT tissue following PKG activation. This upregulation may serve to constrain overactivation of 5-HT_1B_ autoreceptors that could detrimentally limit social engagement. While we do not understand the basis of the sex-specificity of elevated SERT surface expression, we are reminded that DAT proteins exhibit sex-specific trafficking in response to presynaptic autoreceptor activation[63], suggesting conserved, yet poorly understood, mechanisms that link transporter trafficking to neurotransmitter release.

Given the central role of somatodendritic 5-HT_1A_ autoreceptors in regulating DRN^5-HT^ neuron excitability, we evaluated whether enhanced 5-HT_1A_ autoreceptor activity is responsible for reduced serotonergic output that fails to sustain repeated social interactions in Ala276 females. Treatment of animals with a full 5-HT_1A_ antagonist only partially rescued sociability in Ala276 females, possibly reflecting equivalent targeting of somatodendritic autoreceptors and postsynaptic 5-HT_1A_ heteroreceptors whose behavioral influences can oppose one another[64]. Prior research has shown that low doses of the partial 5-HT_1A_ agonist buspirone preferentially targets DRN 5-HT_1A_ autoreceptors[42-44] when given systemically. Indeed, we found that buspirone restored social preference and partially restored NAc extracellular 5-HT responses during repeated social interactions in Ala276 females. In contrast, buspirone reduced sociability and 5-HT signaling in WT females, consistent with sufficient 5-HT clearance by WT SERT to shield 5-HT_1A_ autoreceptors and limit inhibition. Interestingly, we found that buspirone impacted bouts of interaction more than their duration. This dissociation suggests that these components may rely on distinct neural signals and/or circuits with initiation of social interaction more sensitive to DRN^5-HT^ activation, whereas local control of 5-HT release and response may figure more prominently in duration of interactions. Although postsynaptic 5-HT_1A_ receptors can indirectly influence serotonergic circuit activity through local feedback networks, the restoration of DRN serotonergic neuron firing by WAY-100635 in acute slices supports a primary role for somatodendritic 5-HT_1A_ autoreceptors in mediating the Ala276 social phenotype. Furthermore, our data support a model whereby buspirone stimulates 5-HT_1A_ autoreceptors, leading to inhibition of DRN^5-HT^ excitation in WT mice. However, impaired activity-dependent upregulation of SERT in Ala276 mutants may elevate somatodendritic extracellular 5-HT, creating conditions under which the functional antagonist properties of buspirone could predominate. Our findings are consistent with the reported benefits of buspirone in reducing social impairment in ASD patients[44] and raise the possibility that overstimulation of 5-HT_1A_ autoreceptors for some individuals may contribute to social behavior deficits associated with ASD. Additionally, buspirone is FDA approved for the treatment of generalized anxiety disorder (GAD) and used off label for the treatment of depression, or depression with GAD, though not without significant side effects, particularly at higher doses[65]. The altered social interactions of these disorders constitutes an important dimension of their presentation[66] and may be sensitive to balancing of SERT-mediated autoreceptor regulation emphasizing the translational relevance of our findings [67].

Complementing the alterations in somatodendritic serotonergic regulation, our findings provide evidence for sex-specific changes in postsynaptic 5-HT receptor signaling in Ala276 mice. Administration of DOI, a 5-HT_2A_ receptor agonist whose head-twitch response depends predominantly on cortical 5-HT_2A_ receptor activation[68], produced a significantly reduced head-twitch response in Ala276 females relative to WT littermates, whereas males were unaffected (Fig. S3G,H). Whether this reduced response reflects changes in receptor expression, signaling efficacy, or compensatory adaptation to altered serotonergic tone remains to be investigated. Regardless, these findings are likely paralleled by changes in 5-HT receptor stimulation and associated behaviors in other brain circuits receiving serotonergic innervation and await further evaluation.

Our results indicate that SERT phosphoregulation supports social function in both sexes, but along distinct behavioral domains: dominance in males versus sociability in females. 5-HT signaling has been linked to the establishment of social hierarchies [69-72]. We found in our earlier study[34] that male, but not female Ala276 mice, showed reduced wins in the tube test, a result indicating diminished social dominance[69]. Pharmacological engagement of 5-HT_1A_ receptors with buspirone normalized the reduced win rate observed in Ala276 males, paralleling its ability to rescue sociability deficits in females. Notably, this rescue occurred despite the absence of a significant genotype difference in NAc 5-HT levels in males, indicating that dominance-related phenotypes may be mediated by serotonergic signaling in circuits distinct from those governing sociability. Candidates include DRN^5-HT^ projections to regions implicated in aggression and social hierarchy such as the medial prefrontal cortex[70], hypothalamus[71], and amygdala[72]. However, lesions or chemogenetic manipulation within NAc shell/core MSNs bidirectionally shift tube-test dominance in male mice, establishing a direct role for NAc circuitry in hierarchy expression[73]. Dominant males also show higher oxytocin receptor binding in the NAc core than subordinates, and intra-NAc shell oxytocin microinjection reduces tube-test dominance, establishing a causal role for oxytocin in rank expression in the NAc[74, 75]. Thus, the lack of alterations in social stimulus elicited 5-HT release in Ala276 mice may reflect NAc subregion-selective 5-HT dynamics. Evaluation of 5-HT elevations within both the NAc core and shell across sexes and genotypes may shed light on these complex circuit-behavior relationships.

Although phosphorylation-dependent regulation of monoamine transporters has long been recognized[25], most evidence derives from heterologous systems, limiting insight into the impact of SERT regulation to behavioral outcomes. The Ala276 model provides the first test of constitutive loss of a PKG-sensitive SERT phosphorylation site *in vivo*, revealing that the capacity for SERT phosphorylation contributes to sociability and to sustaining NAc 5-HT signaling during repeated social interactions, particularly in females. Although our studies were conducted in mice, these findings may have implications for the actions of psychoactive drugs that target 5-HT signaling. For example, MDMA, a molecule that triggers 5-HT efflux via SERT has been shown to promote social engagement[17]. While addiction liability and safety concerns constrain its use in disorders that feature social deficits, social effects of the drug are mediated by NAc 5-HT receptors signaling[8, 12]. Additionally, SSRIs that also elevate extracellular 5-HT have been prescribed for ASD treatment in children and adults, but evidence for reductions in the core symptoms of ASD including social function by SSRIs are lacking[76]. Our data suggest that targeting SERT regulation, either upstream via the nNOS-cGMP-PKG pathway or via selective modulation of 5-HT_1A_ autoreceptors, may provide a more precise strategy to improve sociability. Unlike SSRIs, which tonically inhibit SERT independent of behavioral state, phosphorylation-dependent regulation of SERT may permit rapid, activity-dependent enhancement of transporter function selectively during periods of elevated DRN^5-HT^ firing. Such mechanisms may therefore provide a fundamentally different therapeutic strategy for disorders in which flexible serotonergic regulation is compromised. Indeed, disrupting the SERT-nNOS interaction was recently proposed as a fast-onset antidepressant mechanism[77]. Evaluation of SERT-nNOS interaction in the context of social behavior awaits testing. Interestingly, these interactions were shown to occur at somatodendritic sites, but not in serotonergic axons, revealing a further complexity in SERT regulation that is only beginning to be assessed. Future work should also seek to disentangle pre-versus postsynaptic 5-HT_1A_ contributions (e.g., site-specific microinfusions), test estrous-stage influences on female phenotypes, and leverage circuit specific modulation to isolate pathways that dictate sex-specific behavioral reliance on SERT phosphorylation.

In summary, our study is the first to identify the capacity to phosphorylate SERT as a molecular mechanism linking serotonergic output to social behavior. By revealing a sex-specific association between enhanced 5-HT_1A_ autoreceptor-dependent inhibition, impaired sustained NAc 5-HT levels, and social dysfunction, and by demonstrating pharmacological rescue with buspirone, we provide mechanistic insight and a potential translational path toward treating social deficits, a core unmet symptom domain in ASD. Given the established contribution of 5-HT_1A_ autoreceptor-mediated inhibition to delayed SSRI response[78-80], our findings suggest that strategies aimed at enhancing SERT Thr276 phosphorylation may offer novel therapeutic opportunities for depression and other disorders treated with SSRIs, while also providing insight into their underlying pathophysiology.

## ACKNOWLEDGEMENTS

We thank Rania Katamish for training in biochemical methods, and Felix Mayer for instruction in acute brain slice preparation and fiber photometry used in this study. We are grateful to Ashley Gordon and the Florida Atlantic University Comparative Medicine staff for expert animal care and husbandry. We also thank Anna Stauffer for exemplary laboratory administration. Behavioral testing was supported by the FAU Stiles-Nicholson Brain Institute Neurobehavior Core, overseen by Maureen Hahn. This work was supported by the NIH grant MH094527 (to R.D.B).

## AUTHOR CONTRIBUTIONS

C. A. and R.D.B. conceived the study, with essential input from A.S. and S.R. C.A. performed all behavioral experiments and the majority of biochemical experiments. A.E. contributed to biochemical experiments, including SDS-PAGE and Western blot analyses. A.S. and T.C.W. assisted with synaptosomal assays, stereotaxic surgeries, and fiber photometry recordings. H.I. performed and analyzed electrophysiological recordings. M.G.Z. carried out histological experiments and assisted in maintaining the mouse colony. C.A. wrote the manuscript with input and editorial revisions from R.D.B., A.S., and S.R. All authors contributed to data interpretation and manuscript preparation.

## MATERIALS AND METHODS

### Animals

Wild type (WT) and homozygous Ala276 littermate mice, between 8-16 weeks of age, were bred from heterozygous breeders maintained on a C57BL/6J background[34]. Mouse genotyping was performed by Transnetyx, Inc. (Cordova, TN, USA). Animals were housed on a 12:12 light/dark cycle (lights on/off at 6am and 6pm, respectively). Experiments were performed during the inactive (light) phase, and mice were housed with 2-5 mice/cage and allowed access to standard chow and water *ad libitum*. All animal experiments were performed under a protocol approved by the Institutional Animal Care and Use Committee (IACUC) at Florida Atlantic University.

### Drugs

Paroxetine hydrochloride (CAS 78246-49-8, cat. no. ab120069, Abcam, Cambridge, MA, USA; 10 µM) was dissolved in synaptosomal assay buffer. WAY-100635 maleate salt (CAS 634908-75-1, cat. no. W108, Sigma-Aldrich, St. Louis, MO, USA; 0.1 mg/kg), buspirone hydrochloride (CAS 33386-08-2, cat. no. B190, Sigma-Aldrich, St. Louis, MO, USA; 0.1 mg/kg), and 8-hydroxy-2-(di-n-propylamino)tetralin hydrobromide (8-OH-DPAT; CAS 87394-87-4, cat. no. H140, Sigma-Aldrich, St. Louis, MO, USA; 1 mg/kg) were dissolved in sterile saline for subcutaneous (s.c) or intraperitoneal (i.p.) injection. 8-Bromo-cGMP (CAS 51116-01-9, cat. no. B1381, Sigma-Aldrich, St. Louis, MO, USA; 250 µM) was dissolved in artificial cerebrospinal fluid (aCSF) for *ex vivo* treatment. d,l-MDMA hydrochloride (CAS 64057-70-1, cat. no. M-032, Lipomed, Cambridge, MA, USA; 7.5 mg/kg) was dissolved in saline for i.p. injection.

### *In vivo* Fiber Photometry

8-9-week-old mice were anesthetized using isoflurane (5% induction/1-2% maintenance) and their heads were fixed in a stereotaxic frame (Harvard Apparatus, Holliston, MA, USA). Ophthalmic ointment was applied to prevent drying of the eyes. 50 µL of sterile saline containing 0.05% bupivacaine and 0.2% lidocaine was administered locally and ketoprofen (10 mg/kg, s.c.) was administered systemically. Following a midline incision, lambda and bregma were levelled and three 1.6 mm screws (Fisher Scientific, Waltham, MA, USA) were affixed to the skull to stabilize the probe. A craniotomy was performed with a Dremel drill allowing for viral injection and probe placement in the NAc (0.8 AP, 1.5 ML and -4.3 DV from bregma). A 34-gauge metal needle attached to 10 μL Nanofil Hamilton syringe (Hamilton, Reno, NV, USA) was carefully lowered into the NAc using the coordinates given above and held in place for 5 min. Next, the needle was slowly withdrawn to -4.2 DV and held in place for 3 additional min. 1 µL containing 1 x 10^10^ genome copies of AAV-hSyn-5-HT_3.0_ (GRAB_5HT3.6_; WZ Biosciences, Columbia, MD, USA) was then injected at an infusion rate of 100 nL per min. After injection, the needle was held in place for 5 min, raised to -4.0 DV, held for an additional 3 min, and slowly withdrawn. Optic probes were constructed by gluing 400 μm core/425 μm cladding diameter optical fiber (Thorlabs, Newton, NJ, USA) to 1.25 mm metal ferrules (Doric Lenses, Quebec, QC, Canada) and polished prior to implantation in the NAc. Probes were affixed to the skull and incisions sealed with dental acrylic. Mice were singly housed following surgery and recordings proceeded at least 21 days after surgery to allow for sensor expression.

To monitor signal from the GRAB_5-HT_ sensor, a light-emitting diode (465 nm, Doric Lenses, Quebec City, QC, Canada) was attached to the optic probe via a zirconia mating sleeve (Doric Lenses) coated black to minimize light escape. Emitted light was band-pass filtered (460-490nm) and detected by a Newport Femtowatt silicone PIN photodetector (New Focus, San Jose, CA, USA). Fiber photometry data were acquired with Synapse Software (Tucker-Davis Technologies, Alachua, FL, USA) controlling an RZ5P lock-in amplifier (Tucker-Davis Technologies). 465 nm excitation was delivered at 210Hz by an LED driver (Doric lenses) in low power mode. A demodulated signal was low pass filtered at 6 Hz and digitized at 1017 Hz. Data were extracted to .csv files using OpenBrowser software (OpenEx, Tucker-Davis Technologies) and analyzed in Excel (Microsoft Corporation, Redmond, WA, USA). Data compilation, visualization, and statistical analysis were performed with Prism 10 (GraphPad Inc., La Jolla, CA, USA). We utilized built-in Excel formulas to determine signal peak, peak frequency, and time to peak.

### Behavioral Assays

Behavioral testing was completed in the Neurobehavior Core Facility under the auspices of the Florida Atlantic University Stiles-Nicholson Brain Institute. Mice were allowed to habituate to testing rooms for 20 min prior to the experiment start. Each experiment combines data from at least two independent cohorts wherein experimenters were blinded to animal genotype.

### Social Preference

The three-chamber preference test was performed as described in the standardized social preference protocol for measuring social deficits in mouse models of autism[81]. The three-chamber apparatus was an acrylic box with clear walls and two outer chambers (18×23cm) connected by a center chamber (18×23cm). Translucent acrylic walls have a 10cm gap for mouse passage with the possibility to insert acrylic walls to separate chambers fully. An inverted wire pencil cup made of steel was used to house the social and non-social stimuli. 150ml glass bottles filled with desiccant were placed on top of each pencil cup to prevent test mice from climbing. All social stimuli mice were habituated for two consecutive days to the pencil cups for 1 hr each. All mice were habituated to the apparatus containing empty pencil cups in the outer chambers, allowing them to explore freely for 10 min. Mice were briefly returned to their home cage while two identical paper balls were placed in the center under each inverted pencil cup for the 10 min pre-test phase. An unfamiliar age-, sex- and strain-matched WT mouse was placed under one of the inverted pencil cup. The placement of the social stimulus mouse was alternated in each round. A lid of a 1L Pyrex glass bottle was used as an unfamiliar, inanimate object that was placed under the other pencil cup as the non-social stimulus. The experimental mouse was allowed to freely explore all three chambers for 10 min. At the end of all the test session, mice were returned to their respective home cages and the apparatus including all pencil cups and glass bottles were cleaned with 70% ethanol to remove any odors for subsequent tests. For studies using pharmacological intervention, mice were injected 10 min prior to the habituation test with either vehicle or 0.1mg/kg WAY 100635 to allow for 30 min drug treatment before the social preference test. Mice were injected with vehicle or 0.1mg/kg buspirone i.p. 30 min prior to habituation test to allow for 50 min drug treatment before exposure to a stimulus mouse [45].

For all test session videos were acquired by a ceiling-mounted digital camera and analyzed using the animal-tracking software EthoVision XT (Noldus Information Technology, Wageningen, The Netherlands). On each experimental day, fidelity of automated mouse tracking was verified. Social investigation in the sniffing zone was quantified as the cumulative time the experimental mouse’s nose point was detected within a predefined 1.5-cm-radius zone surrounding each wire pencil cup. A sociability index was calculated[12] as 100 x (time spent with social stimulus - time spent with non-social stimulus)/(time spent with social stimulus + time spent with non-social stimulus). Mice that failed to move 1000cm in 10 min in exploration test or failed to explore one of the cups for more than 10s was excluded from further analysis.

### Juvenile Interaction

To assess 5-HT dynamics in the NAc, we introduced a novel juvenile (4–5 weeks of age), sex-matched conspecific into the home cage of WT or Ala276 mice with implanted optic probes for monitoring of social stimulus-dependent 5-HT release. We utilized juvenile stimulus mice for these experiments to minimize territorial aggression and encourage social exploration. Stimulus mice were introduced at the opposite end of the cage from the experimental mouse and remained in the cage for a 5-min recording period. Experiments were video recorded and later scored by a blinded observer for the frequency and duration of active interaction bouts, which required orienting of the experimental subject toward the stimulus mouse and included sniffing, following, and/or grooming behaviors. A social interaction ratio was calculated as the percentage of total interaction time during which the experimental mouse was actively engaged (active interaction time/total interaction time × 100), with total interaction time including interactions initiated by either mouse. This measure was used to distinguish active social engagement from passive participation in social interactions. Behavioral annotations were aligned to photometry traces to match 5-HT release dynamics to instances of active interaction.

### DOI Induced Head Twitch

DOI-induced head twitch was conducted as previously described[82]. Briefly, Ala276 mice and their SERT Thr276 littermates treated with either saline or DOI (1.0 mg/kg, i.p.; Sigma-Aldrich, St. Louis, MO, USA) and placed in a new cage without a food tray or lid to ensure proper video recording using a go-pro camera (GoPro Inc., San Mateo, CA, USA). Mice were video recorded for 49 min. Head twitches over a period of 15 min after 34 min post-treatment were counted by two independent reviewers blinded to genotype. Head twitches were quantified and analyzed.

### Light/Dark Transition Test

Anxiety-like behavior was determined by exploiting the conflict in mice between spontaneous exploration and their aversion to brightly illuminated spaces[83]. An insert divided Med Associates locomotor chambers into 2 equally sized spaces, one brightly illuminated and one dark, linked by a door. Animals were placed in the dark chamber and allowed to freely explore both chambers for 10 min. The latency to return to the light side, total time spent on the light side and total distance traveled were recorded for each animal.

### Evaluation of SERT Surface Trafficking in Acute Brain Slices

Brain slice preparation and experiments were performed under constant oxygenation (95%O2:5%CO2). Animals were sacrificed by rapid decapitation and excised brains were moved to ice-cold sucrose-artificial CSF (S-aCSF; in mM: sucrose 250, KCl 2.5, NaH2PO4 1.2, NaHCO3 26, D-glucose 11, MgCl_2_ 1.2, CaCl_2_ 2.4, pH 7.4, 300-310mOsm). A vibratome (Leica VT100S; Leica Microsystems, Wetzlar, Germany) was used to prepare 290µm slices that contained the DRN, the dStr and vStr(matching coordinates of 0.2-0.5mm AP for dStr, and 1.0-1.5mm AP for vStr in the Allen Mouse Brain Atlas) in S-ACSF following which, slices were allowed to recover at 30-32°C for 1h in ACSF, substituting 92mM NaCl for sucrose (pH 7.4, 300-310mOsm). Before drug treatment, slices were washed with ACSF at 37°C and then treated with vehicle or 8-Bromo-cGMP (CAS 51116-01-9, cat. no. B1381, Sigma-Aldrich, St. Louis, MO, USA; 250 µM) for 30min. Slices were washed in ice-cold ACSF, followed by treatment with sulfo-NHS-SS-biotin (1 mg/ml, Thermo Fisher Scientific, Waltham, MA, USA) on ice for 30 min. Reactions were quenched by treatment with 0.1 M glycine two times for 10 min, followed by rapid washes with ice-cold ACSF. dStr, vStr, and DRN were rapidly dissected from whole striatum and midbrain slices and broken up in RIPA (in mM: NaCl 150, KCl 2.5 m, Tris 50, and 1% Triton X-100) .5% sodium deoxycholate and 0.1% sodium dodecyl sulfate (SDS); Sigma-Aldrich, St Louis, MO) containing a protease inhibitor cocktail (Sigma-Aldrich, St Louis, MO, USA) using a 29g insulin syringe. After 50 min of lysis under rotation at 4°C, protein concentrations were determined using the BCA protein assay (Thermo Fisher Scientific, Waltham, MA, USA) with bovine serum albumin (BSA; Thermo Fisher Scientific, Waltham, MA, USA) as standard, with a FLUOstar Omega microplate reader (BMG Labtech, Ortenberg, Germany). Isolation of biotinylated proteins was performed as described by Gabriel et al.[84]. Briefly, equal amounts of protein for each sample were added to washed streptavidin agarose beads (Thermo Fisher Scientific, Waltham, MA, USA) at a ratio of 20µg protein per 50µl bead slurry and mixed overnight at 4°C. Following washes with lysis buffer, protein was eluted/denatured by adding 2X Laemmli buffer (Bio-Rad Laboratories, Hercules, CA, USA) at room temp, shaking for 30 min. After pelleting the beads, all samples were subjected to SDS-PAGE (10% Gels, Thermo Fisher Scientific, Waltham, MA, USA). Following SDS-PAGE, proteins were transferred to PVDF membranes (MilliporeSigma, Burlington, MA, USA) at 4°C for 18h at 20V and then blocked with 5% dry milk in room temp wash buffer (in mM: NaCl150, 2.5 KCl, 50 Tris, and 0.1% Tween 20). Blocked membranes were incubated with guinea-pig anti-SERT (HTT-GP-Af1400, Nittobo Medical, Tokyo, Japan) and rabbit anti-TH (2025-THRAB, PhosphoSolutions, Aurora, CO, USA; RRID:AB_2492276) solution of 1:1000 in wash buffer overnight at 4°C. Membranes were then rinsed 4 in wash buffer before incubation with goat anti-guinea HRP-conjugated secondary antibody at a 1:10,000 dilution for 1h. Following multiple washes using wash buffer, membranes were subjected to chemiluminescent visualization (Bio-Rad Laboratories, Hercules, CA, USA). Immunoreactivity was detected using the Odyssey Fc (LI-COR Biosciences) followed by densitometry analysis of protein levels normalized to actin using ImageJ. Blot images were minimally processed during quantitation and cropped for the presentation of representative figures.

### Synaptosomal Transport Assays

Midbrain and striatal SERT activity was assessed using synaptosomes. Mice were sacrificed by rapid decapitation followed by dissection of the midbrain and striatum on ice. Tissue was homogenized in ice-cold sucrose (0.32 M; Sigma-Aldrich, St. Louis, MO, USA). After a 10 min, 4°C centrifugation at 800xg, supernatants were transferred to fresh tubes followed by centrifugation at 15,000 xg at 4°C for 20 min. Synaptosomal pellets were resuspended in Krebs-Ringers-HEPES (KRH) assay buffer (in mM: NaCl 130, KCl 1.3, CaCl_2_ 2.2, MgSO_4_ 1.2, KH_2_PO_4_ 1.2, HEPES 10, glucose 10; pH 7.4) supplemented with pargyline (CAS 555-57-9, cat. no. P8013, Sigma-Aldrich, St. Louis, MO, USA; 100 µM) and ascorbic acid (CAS 50-81-7, cat. no. A92902, Sigma-Aldrich, St. Louis, MO, USA; 100 µM). After assessing protein concentrations using the BCA method (ThermoFisher, Waltham, MA), 50 μg synaptosomal protein per tube were incubated for 10 min at 37°C with MDMA 10^−9^-10^−4^ M prior to the addition of 50 nM [3H]5-HT (specific activity 28 Ci/mmol; PerkinElmer, Waltham, MA). Uptake was terminated after 10 minutes by rapid washing with ice-cold PBS onto Whatman filters pre-soaked in 0.3% polyethyleneimine (CAS 9002-98-6, Sigma-Aldrich, St. Louis, MO, USA) using a Brandel Cell Harvester (Brandel, Gaithersburg, MD USA). Captured [^3^H]5-HT was eluted in scintillation liquid (Ecoscint H, National Diagnostics, Charlotte, NC, USA) overnight and quantified using a Wallac scintillation counter (PerkinElmer, Waltham, MA, USA). Specific uptake was calculated by subtracting the counts obtained from samples incubated in parallel with 10 μM paroxetine from total counts without paroxetine.

### Whole Cell Patch Clamp Recordings in Acute Brain Slices

Coronal slices (280 µm) containing the DRN were prepared using a vibratome (VT1000S, Leica Biosystems, Wetzlar, Germany) in ice-cold, carbogenated (95% O_2_/5% CO_2_) sucrose-substituted artificial cerebrospinal fluid (aCSF; in mM: NaCl 87, sucrose 75, KCl 2.5, NaH_2_PO_4_ 1.25, glucose 25, MgCl_2_ 7, CaCl_2_ 0.5, NaHCO_3_ 25 ). Slices recovered for 30 min in the same solution at 28 °C and were subsequently equilibrated for 1 h at room temperature in standard aCSF (in mM: NaCl 124, KCl 2.5, NaH_2_PO_4_ 1, CaCl_2_ 2.5, MgSO_4_ 1.3, NaHCO_3_ 26, glucose 20; 310 mOsm/kg). During recordings, slices were submerged in a chamber (RC-27, Warner Instruments, Hamden, CT, USA) mounted on an upright microscope (Axioskop 2 FS, Zeiss, Oberkochen, Germany) equipped with infrared differential interference contrast (IR-DIC) optics. Images were acquired through a 40× water-immersion objective using a cooled CCD camera (ORCA-R2, Hamamatsu Photonics, Hamamatsu, Japan) controlled by HCImage software (Hamamatsu Photonics). Slices were continuously perfused with carbogenated aCSF via a peristaltic pump and in-line heater (TC-324B, Warner Instruments) at 28 °C, with complete exchange of chamber volume in ∼3 min. Recording electrodes (5-7 MΩ) were pulled from borosilicate glass capillaries (cat. no. 8250, A-M Systems, Sequim, WA, USA) using a micropipette puller (P-97, Sutter Instrument, Novato, CA, USA). The internal pipette solution contained (in mM): K-gluconate 135, KCl 10, HEPES 10, EGTA 1, MgATP 1, Na_2_GTP 0.2, and Na_2_-phosphocreatine 1 (pH 7.3, adjusted with KOH; 295 mOsm/kg). Whole-cell current-clamp recordings were obtained with a MultiClamp 700B amplifier (Molecular Devices, San Jose, CA, USA). Signals were low-pass filtered at 2 kHz, digitized at 5 kHz with an Axon Digidata 1550B (Molecular Devices), and acquired using pClamp 10.7 (Molecular Devices). Series resistance was monitored throughout but not compensated; recordings with access resistance >20 MΩ were excluded. Putative serotonergic neurons were identified according to previously described electrophysiological criteria[85], including a gradual initial phase of the afterhyperpolarization. To assess autoreceptor regulation, neurons were activated with the α_1_-adrenergic agonist phenylephrine (CAS 59-42-7, cat. no. P6126, Sigma-Aldrich, St. Louis, MO, USA; 10 µM), and maximum firing rates at I = 0 were measured under basal conditions and following perfusion with the 5-HT_1A_ receptor antagonist WAY-100635 (CAS 634908-75-1, cat. no. W108, Sigma-Aldrich, St. Louis, MO, USA; 50–100 nM). Data were analyzed using Clampfit 10.7 (Molecular Devices), OriginPro 2017 (OriginLab, Northampton, MA, USA), and Prism 10.0 (GraphPad Software, La Jolla, CA, USA).

### Statistical Analyses

All statistical analyses were performed using Prism 10 (GraphPad Software, La Jolla, CA, USA). Data are presented as mean ± S.E.M., and statistical significance was defined as *P* < 0.05. Comparisons between two independent groups were conducted using unpaired, two-tailed Student’s *t* tests. Experiments involving two independent variables were analyzed using two-way analysis of variance (ANOVA), followed by Sidak’s or Fisher’s LSD post hoc tests. Repeated-measures ANOVA was used for analyses of fiber photometry traces across social interaction sessions to account for within-subject variability over time.

