## Supplementary Data for "Social Behavior in Mice Relies on a Capacity to Phosphorylate the Serotonin Transporter"

### SUPPLEMENT

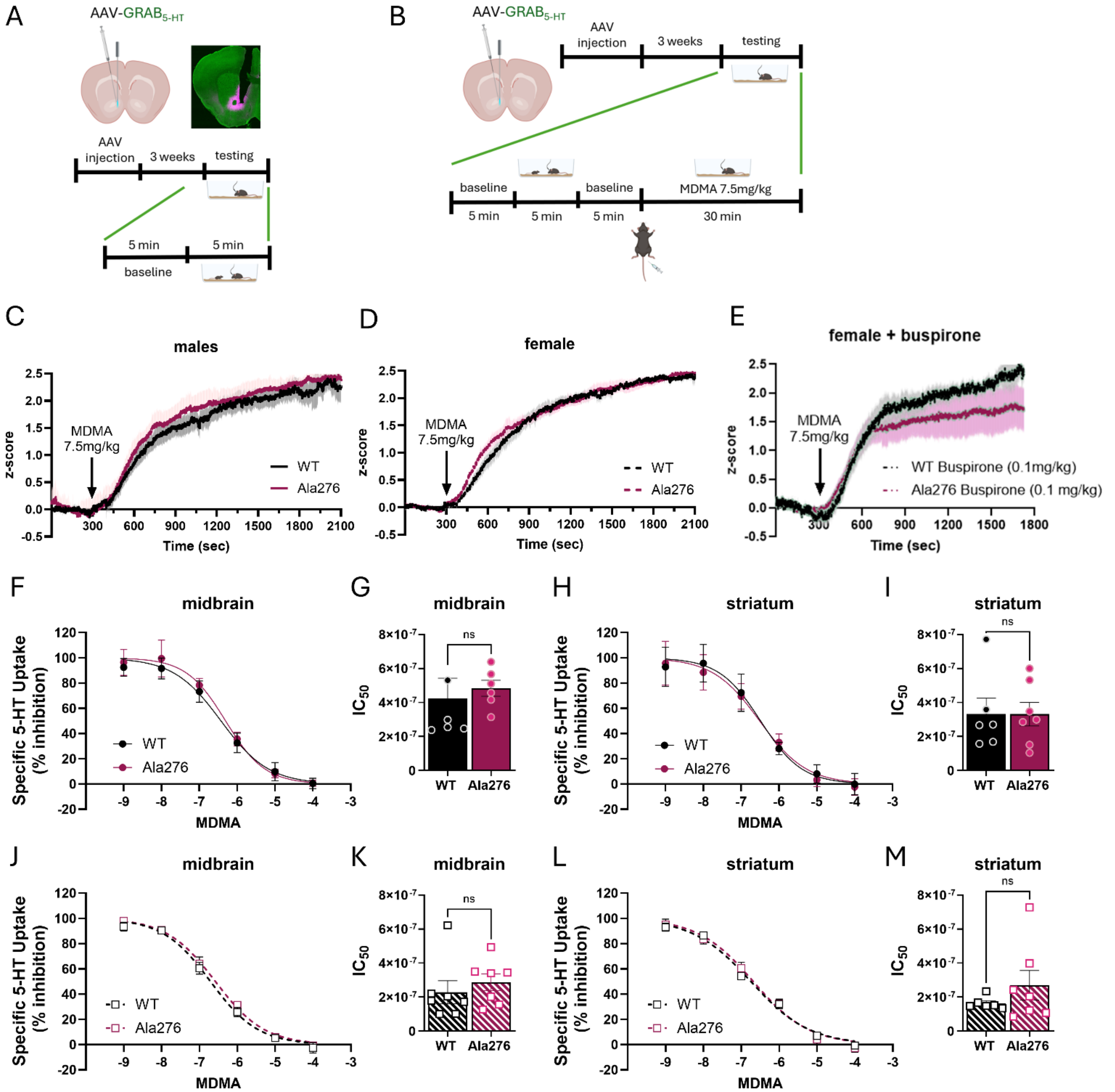


**Figure S1. MDMA-evoked NAc 5-HT signals and MDMA inhibition of synaptosomal uptake are intact in Ala276 mice.** (**A/B**) Schematic of optic-fiber placement in the nucleus accumbens (NAc) medial shell and timeline. $GRAB_{5\text{-}HT}$ was expressed via AAV and fluorescence was monitored by fiber photometry. (**C-E**) Averaged $GRAB_{5\text{-}HT}$ fluorescence traces (NAc) during MDMA challenge in males (WT n=7, Ala276 n=8) (**C**), females (WT n=7, Ala276 n=6) (**D**), and females pretreated with buspirone (WT n=8, Ala276 n=7) (**E**). MDMA (7.5 mg/kg, i.p.) was delivered at t = 300 s (arrow). WT and Ala276 (KI) show comparable responses. (**F-I**) Males: MDMA inhibition of [$^3$H]-5-HT uptake in synaptosomes. Midbrain dose-response curves (**F**) and corresponding $IC_{50}$ summary (**G**) show no genotype differences (WT n=5, Ala276 n=6). Striatum dose-response curves (**H**) and $IC_{50}$ summary (**I**) likewise show no genotype effect (WT n=5, Ala276 n=6). (**J-M**) Females: MDMA inhibition of [$^3$H]-5-HT uptake in synaptosomes. Midbrain curves (**J**) and $IC_{50}$ (**K**) show no genotype differences (WT n = 6, KI n = 7). Striatum curves (**L**) and $IC_{50}$ (**M**) also show no genotype effect (WT n = 6, KI n = 7). Data are mean ± SEM. Traces depict averaged fluorescence aligned to injection; MDMA time indicated by arrow **(C-E**). Synaptosomal dose-response curves were fit by nonlinear regression; $IC_{50}$ values (**G, I, K, M**) compared by unpaired, two-sided Student's t-test. ns, not significant.

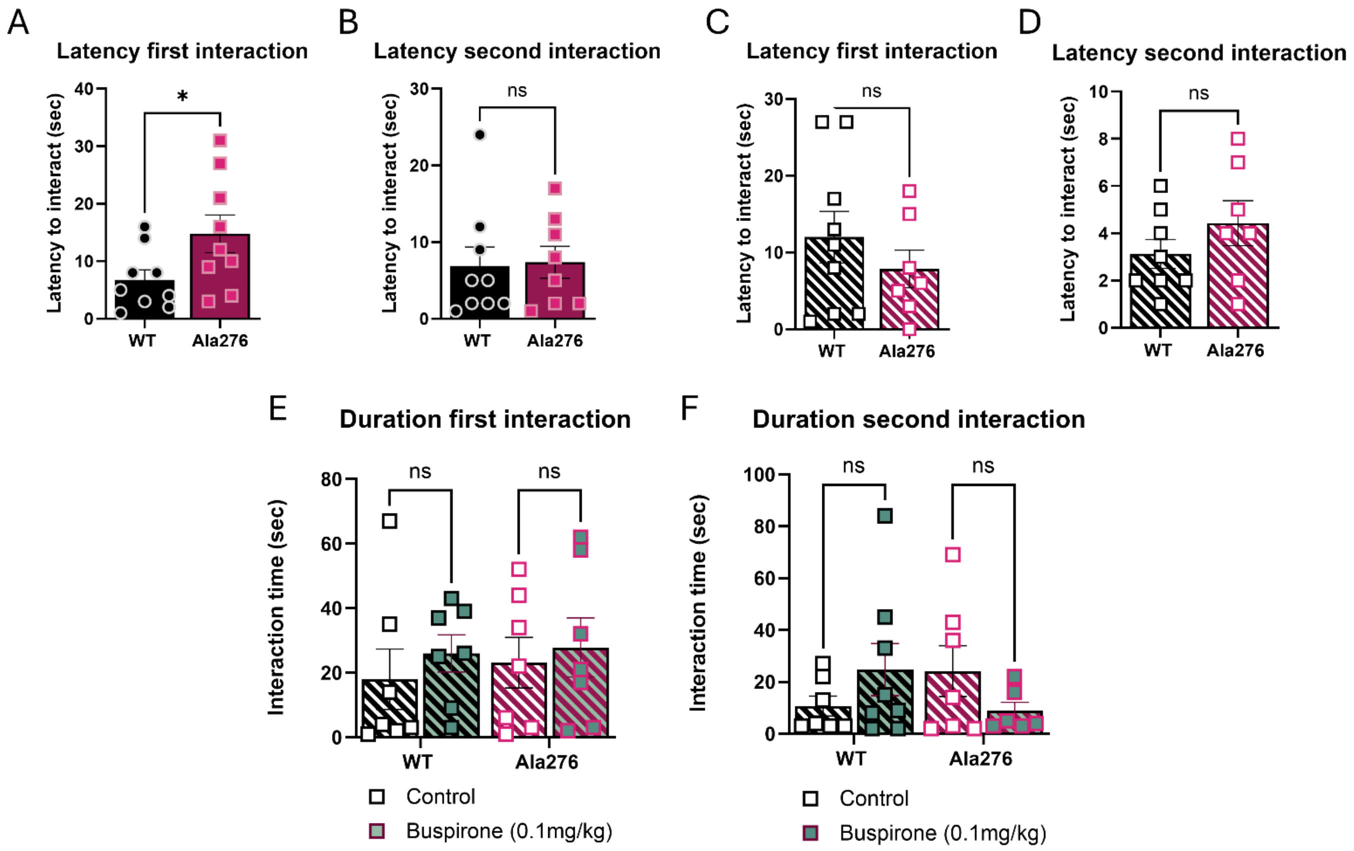


**Figure S2. Juvenile-interaction approach latencies and interaction duration.** (**A-B**) Males (juvenile interaction, fiber photometry cohorts): (**A**) Latency to the first approach is significantly increased in Ala276 compared with WT, indicating a delay in initial social engagement. (**B**) Latency to the second approach does not differ between genotypes. (**C-D**) Females: (**C**) Latency to the first approach and (**D**) latency to the second approach are comparable between WT and KI. (**E-F**) Females ± buspirone. Duration of the first (**E**) and second (**F**) interactions in WT and KI females under vehicle vs buspirone (0.1 mg/kg, i.p., 30 min before test begin) conditions. (**E**) First interaction duration is similar across genotypes and drugs. (**F**) Although not statistically significant, WT mice given buspirone show a modest increase in second-interaction duration, whereas KI mice given buspirone exhibit a shorter second-interaction duration, approximating WT vehicle levels. Data are mean ± SEM. Statistics: unpaired, two-sided Student's t-tests for (A-D); two-way ANOVA (genotype × drug) with Sidak's post hoc tests for (E-F). ns, not significant; *$p < 0.05$.

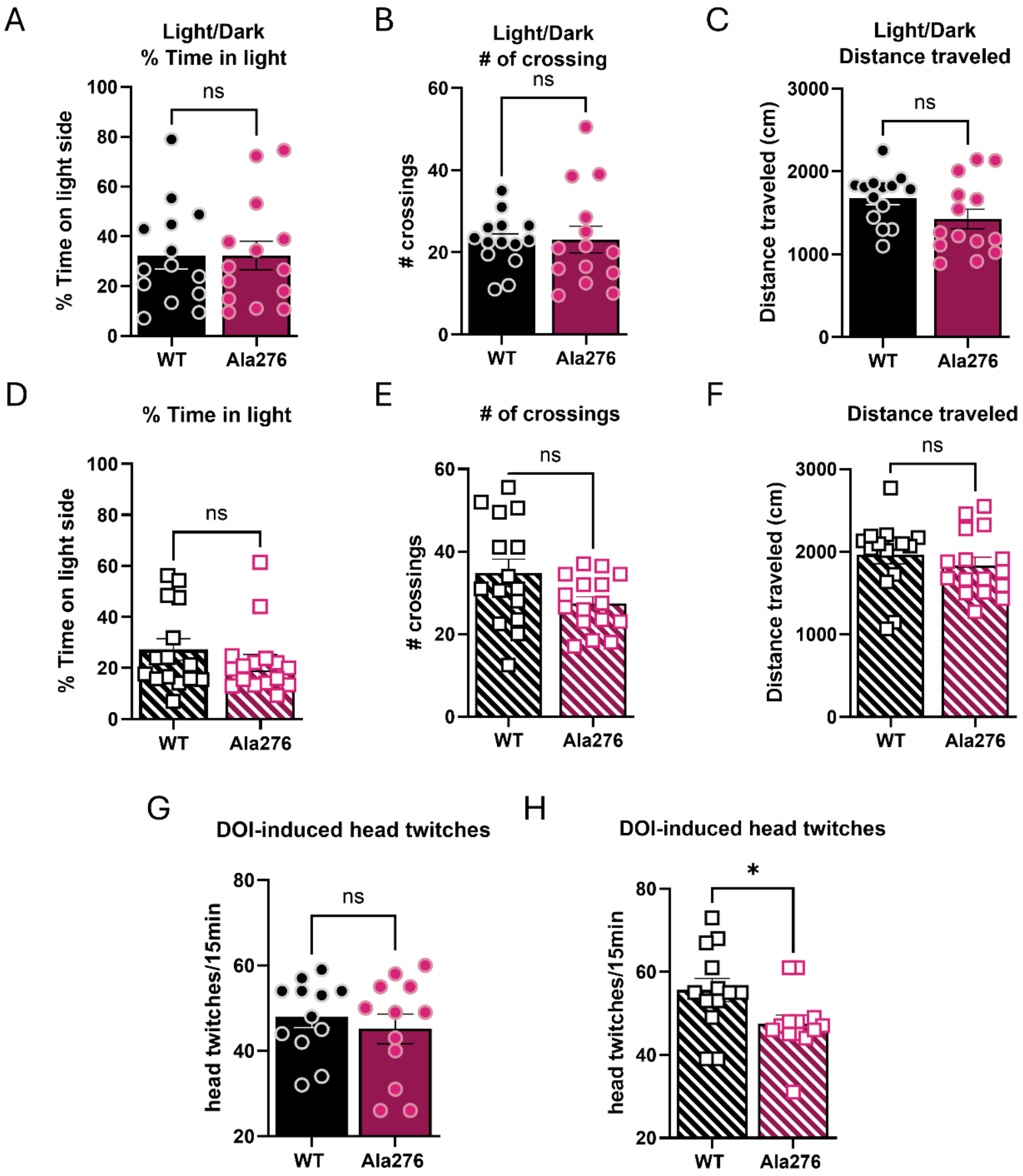


**Figure S3. Ala276 mice do not exhibit anxiety-like behavior while DOI-evoked head-twitch response is reduced in females.** (**A-F**) Light/dark test (10 min), males (n=14): percent time spent in the light compartment (**A**), number of light↔dark crossings (**B**), and total distance traveled (**C**) show no differences between WT and Ala276 mice. (**D-F**) Light/dark test (10 min), females (WT n=15, Ala276 n=16): percent time in light (**D**), crossings (**E**), and total distance (**F**) are comparable between WT and Ala276 mice. (**G-H**) DOI head-twitch response: number of head twitches scored over a 15-min epoch following DOI (1.0 mg/kg, i.p.) in males (**G**; no genotype difference, n=12) and females (**H**; Ala276 significantly reduced vs WT, n=13). Data are presented as mean ± SEM. Statistical comparisons were performed using unpaired, two-sided Student's t-tests. ns, not significant; *$p < 0.05$.

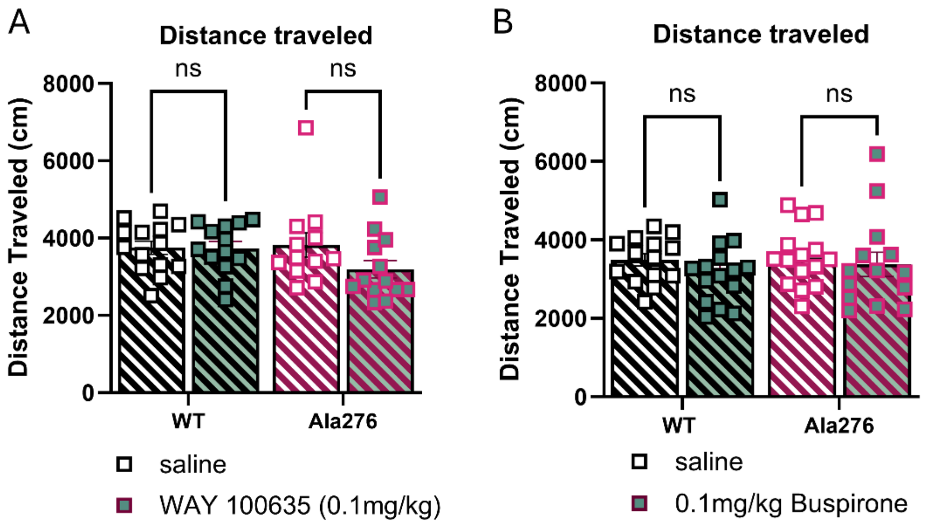


**Figure S4. 5-HT$_{1A}$ receptor manipulations do not affect locomotor activity during the social preference test.** (**A**) Total distance traveled during the 10 min session in WT and Ala276 females pretreated with saline or WAY-100635 (0.1 mg/kg, s.c., 10 min before test begin); no genotype or treatment effects. (**B**) Total distance traveled during the 10 min session in WT and KI females pretreated with saline or buspirone (0.1 mg/kg, i.p., 30 min before test begin); no genotype or treatment effects. Data are presented as mean ± SEM. Statistical comparisons were performed using two-way ANOVA (genotype × treatment). ns, not significant;
